# Oncogenic H-Ras Signaling Differentially Targets Replicative and Specialized DNA Polymerases for Depletion

**DOI:** 10.1101/2020.07.01.180455

**Authors:** Wei-chung Tsao, Raquel Buj, Katherine M. Aird, Julia M. Sidorova, Kristin A. Eckert

**Author notes:** Corresponding author: Kristin Eckert, Penn State College of Medicine, 500 University Drive, H059, Hershey, Pa 17033.

## Abstract

Oncogene activation significantly alters DNA replication dynamics, causing replication stress and genome instability. However, little is known about DNA polymerase expression and regulation during oncogene-induced replication stress. We discovered that the Pol α catalytic subunit, Pol δ, Pol η and Pol κ are all depleted in response to H-Ras^G12V^ overexpression in multiple human cell lines. Distinct transcriptional and post-translational mechanisms mediate replicative and specialized DNA polymerase regulation, respectively, and include both MEK-dependent and -independent pathways. Moreover, Pol η depletion is sufficient to induce a senescence-like growth arrest in non-transformed cells. We provide evidence that H-Ras^G12V^-induced polymerase depletion contributes not only to oncogene-induced replication stress, but also to cell cycle checkpoint enforcement. Polymerase degradation is a protective response, associated with improved cell survival in the face of oncogene-induced stress. Our findings significantly impact our understanding of oncogene-induced cellular transformation and suggest that imbalanced polymerase levels may contribute to neoplastic progression.

## Introduction

DNA replication is a critical phase of the cell cycle that must be tightly regulated to ensure accurate genome duplication. Failure to maintain DNA replication regulation leads to genome instability and ultimately tumorigenesis. Activating mutations in oncogenes alter DNA replication dynamics, promoting replication stress and genome instability (Di Micco et al. 2006; Kotsantis et al. 2018). In turn, cells can undergo proliferative arrest known as oncogene-induced senescence (OIS), which acts as a tumorigenic barrier by preventing neoplastic transformation (Bartkova et al. 2005; Michaloglou et al. 2005; Di Micco et al. 2006; Kotsantis et al. 2018). Replication stress is broadly defined as the slowing or stalling of the replication fork when the replisome encounters obstacles during DNA replication, and is associated with genome instability (Zeman and Cimprich 2014). The replisome is a highly dynamic structure that incorporates multiple DNA polymerases, enzymes that are integral to all replication processes, including ongoing fork elongation, fork restart, and fork repair (Zhu et al. 2005; Lange et al. 2011; Gaillard et al. 2015; Burgers and Kunkel 2017; Lee et al. 2019). The regulation of various DNA polymerases to accomplish replication in response to exogenous, DNA damage-induced replication stress is well understood (Sale et al. 2012; Bournique et al. 2018). Critically, much remains unknown about the impact of oncogenes on DNA polymerase regulation, and the DNA polymerases utilized during endogenous, oncogene-induced replication stress.

Significant evidence supports the hypothesis that the replication stress and DNA damage responses are induced during oncogene activation, prior to senescence (Di Micco et al. 2006; Bartek et al. 2007; Liu et al. 2018). Oncogenic Ras activation causes replicative stress, DNA damage, and OIS through a variety of mechanisms (Bartkova et al. 2006; Di Micco et al. 2006; Aird et al. 2013; Kotsantis et al. 2018). Constitutively activated mutant H-Ras (hereafter referred to as Ras^G12V^) increases CDK2 activity and subsequent G1/S checkpoint abrogation, leading to increased origin firing, hyper-replication, and aberrant cell proliferation (Di Micco et al. 2006; Pylayeva-Gupta et al. 2011). Consequently, the prolonged presence of Ras^G12V^ activity leads to replication stress through increased production of reactive oxygen species, replication-transcription machinery collisions, and depleted dNTP pools (Bartkova et al. 2006; Mannava et al. 2013; Ogrunc et al. 2014; Kimmelman 2015; Miron et al. 2015; Kotsantis et al. 2016).

The fidelity of genome replication is orchestrated by engaging multiple DNA polymerases (Barnes and Eckert 2017). Replicative polymerases delta (Pol δ) and epsilon (Pol ε) replicate the bulk of eukaryotic genomes under unstressed conditions, and are generally regarded as high fidelity (Lujan et al. 2016). Current models to explain resolution of stalled replication forks invoke specialized polymerases to perform DNA synthesis either at the fork, when replicative polymerases are inhibited (Sale et al. 2012), or post-replicative gap-filling synthesis behind the replication fork (Gao et al. 2017). Specialized polymerases eta (Pol η) and kappa (Pol κ) maintain the fidelity of genome duplication through non-B DNA structures, common fragile sites, and DNA lesions (Betous et al. 2009; Rey et al. 2009; Sale et al. 2012; Barnes et al. 2017; Tsao and Eckert 2018; Tonzi and Huang 2019). Correspondingly, replication stress caused by hydroxyurea, aphidicolin, and chemotherapeutic agents induce the up-regulation of Pol η, allowing cells to complete genome replication (Ceppi et al. 2009; Srivastava et al. 2015; Barnes et al. 2018). Recent research has shed some light on the DNA polymerases required to mitigate oncogenic stress. DNA Pol δ facilitates break-induced replication fork repair and cell cycle progression in cells overexpressing cyclin E (Costantino et al. 2014). In normal human fibroblasts and cancer cells, Pol κ is important for the tolerance of Cyclin E/CDK2-induced DNA replication stress (Yang et al. 2017), while Pol η confers tolerance to Myc-induced replication stress in cancer cells (Kurashima et al. 2018).

Given their vital roles in maintaining genome stability, we sought to understand the regulation of DNA polymerases during oncogene-induced replication stress. We report the astonishing discovery that several DNA polymerases are actively depleted in a temporal manner in response to Ras^G12V^ overexpression in human cells. Moreover, we uncovered distinct mechanisms regulating DNA polymerase expression at the transcriptional and post-translational level for replicative and specialized polymerases, respectively. Finally, we present evidence that DNA polymerase depletion in response to Ras^G12V^ signaling is a protective mechanism, allowing cells to adapt to oncogene-induced replication stress by limiting replication fork progression and enforcing checkpoint activation, which ultimately improves cell survival. Our study shows, for the first time, that the levels of DNA polymerases are variable during oncogenic stress, and we suggest that imbalanced polymerase levels may contribute to oncogene-induced genome instability.

## Results

### Overexpression of oncogenic H-Ras induces differential depletion of DNA polymerases in nontumorigenic human cells

We aimed to elucidate DNA polymerase expression in response to oncogene-induced replication stress, using an established experimental model of Ras^G12V^ overexpression (OE) (Di Micco et al. 2006; Tu et al. 2011). We used hTERT-immortalized BJ5a human fibroblasts, to avoid background replicative senescence of primary fibroblasts (Hayflick and Moorhead 1961; Martin-Ruiz et al. 2004). hTERT cells do exhibit an OIS response, although some populations can bypass senescence in the long-term (Kohsaka et al. 2011; Patel et al. 2016; Bianco et al. 2019). hTERT-BJ5a cells were stably transduced with a constitutively active, Ras^G12V^ expressing retroviral vector or empty vector control (Figure 1—figure supplement 1A). As expected, we observed an induction of several senescent phenotypes over the course of several days in Ras^G12V^ OE cells, including increased senescence-associated (SA)-β-galactosidase staining, decreased laminB1 expression and increased p16 expression, compared to control-infected cells (Figure 1—figure supplement 1B-D). After 8 days of Ras^G12V^ OE, we measured significantly altered cell cycle distributions and increased phosphorylation of Chk2 (Figure 1—figure supplement 2A-B). Together, these data show the expected onset of senescent phenotypes and the increased DNA damage checkpoint response after Ras^G12V^ OE.

Next, we measured the expression of several DNA polymerases genes in Ras^G12V^ and control cells as a function of days following oncogene overexpression and the onset of senescent phenotypes. Strikingly, we observed significant downregulation of replicative polymerase genes, including POLA1, POLD1, POLD3, and POLE1, after Ras^G12V^ OE as early as day 2 post selection, and sustained through day 8 (Figure 1A). In contrast, we found only transient and slight downregulation of specialized polymerase genes POLH and POLK at Day 4, while the DNA repair polymerase POLB gene expression was not affected at any time point measured.

**Figure 1.**
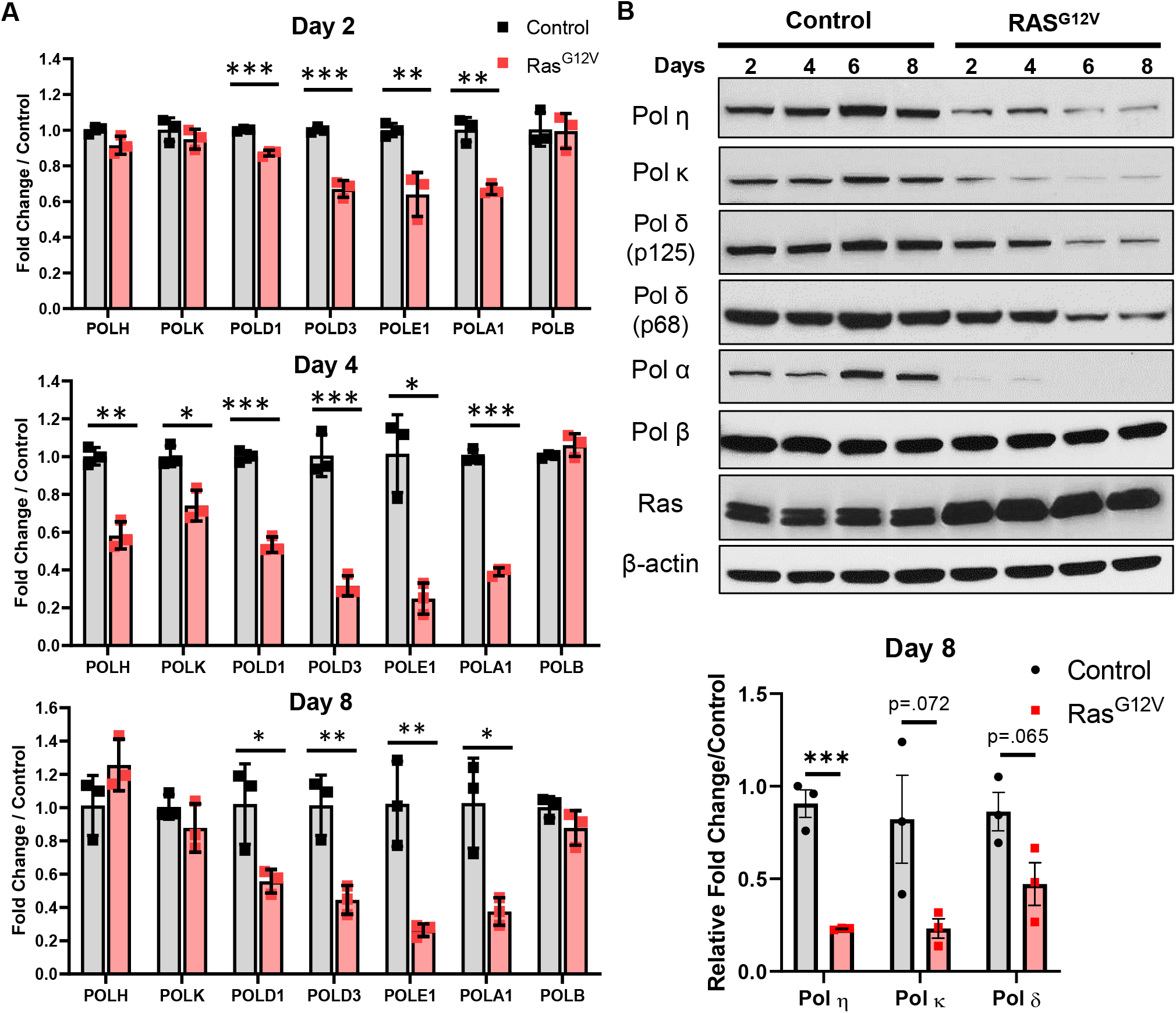
Overexpression of Oncogenic Ras^G12V^ induces downregulation of Replication Fork-Associated DNA polymerases. (A) hTERT BJ5a human fibroblasts were infected with pBabe retrovirus empty vector (control; gray bars) or encoding mutant Ras^G12V^ (red bars). POL gene expression was determined using qRT-PCR at the indicated timepoints after selection. Data represent mean +/− SEM of three biological replicates. (B) *Top panel*-Immunoblot analyses of control and Ras^G12V^ -infected cells at indicated timepoints. One of three biological replicates is shown. *Bottom panel*- Quantification of DNA polymerase protein expression 8 days after control or Ras^G12V^ transduction. Data represent mean +/− SD of three biological replicates. Statistical analyses (A and B) were performed using Holm’s-Sidak multiple t-tests. *p<0.05, **p<0.01, ***p<0.005.

DNA polymerase protein levels were also significantly and differentially impacted by Ras^G12V^ OE (Figure 1B). The catalytic subunit of replicative Pol α was severely depleted as early as day 2. Moreover, expression of Pols η, κ, and both catalytic (p125; POLD1 gene) and accessory (p68; POLD3 gene) Pol δ subunits were depleted in a temporal manner after Ras^G12V^ OE (Figure 1B, Figure 1—figure supplement 1E). However, similar to its mRNA expression, Pol β protein expression is not altered after Ras^G12V^ OE. Taken together, these results show that DNA polymerases are differentially regulated at the transcript and protein levels in response to oncogenic Ras^G12V^ overexpression.

### Ras^G12V^ induced depletion of DNA polymerases is dependent on the proteasomal degradation pathway

We tested whether the depletion of DNA polymerase proteins after Ras^G12V^-induced senescence is due to proteasomal degradation. For each time point after Ras^G12V^ OE, hTERT-BJ5a cells were treated with MG132, a 26S proteasome inhibitor, immediately prior to harvesting cell lysates for analysis. Remarkably, we observed a robust rescue of specialized Pols η and κ at multiple timepoints (Figure 2A and Figure 2-figure supplement 1B) in treated cells expressing Ras^G12V^. Pols η and κ protein levels are also slightly increased after MG132 treatment of control cells, consistent with previous reports (Hendriks et al. 2014; Bertoletti et al. 2017). In contrast, we observed little to no rescue of Pols α (catalytic subunit) or δ (p125 or p68 subunits). This result is consistent with replicative polymerase regulation in response to Ras^G12V^ OE occurring primarily at the transcriptional level (Figure 1A). To establish the generality of this response, we tested an hTERT-immortalized, patient-derived Xeroderma Pigmentosum Variant (hXPV) cell line complemented with an ectopic POLH cDNA vector (hXPVη). Again, Ras^G12V^ OE in the hXPVη cells induced the depletion of Pols η and κ, and this depletion could be rescued by MG132 treatment (Figure 2—figure supplement 1A). Finally, we repeated this experiment using IMR90 primary fibroblasts. As with the hTERT cell lines, Pols η and κ are depleted after Ras^G12V^ OE, and this degradation can be rescued by MG132 treatment (Figure 2B and Figure 2-figure supplement 1B). Notably, we observed a cell type-specific regulation of the Pol δ catalytic (p125) subunit in response to Ras^G12V^ signaling. Specifically, MG132 treatment of Ras^G12V^-infected hXPVη and IMR90 cells partially restored Pol δ (p125) levels, whereas MG132 treatment of BJ5a cells did not (Figure 2B; Figure 2-figure supplement 1B). Together, these data show that during Ras^G12V^ signaling, DNA Pols η, κ and, in some cases, δ (p125) are regulated at the post-translational level through the proteasome degradation pathway.

**Figure 2.**
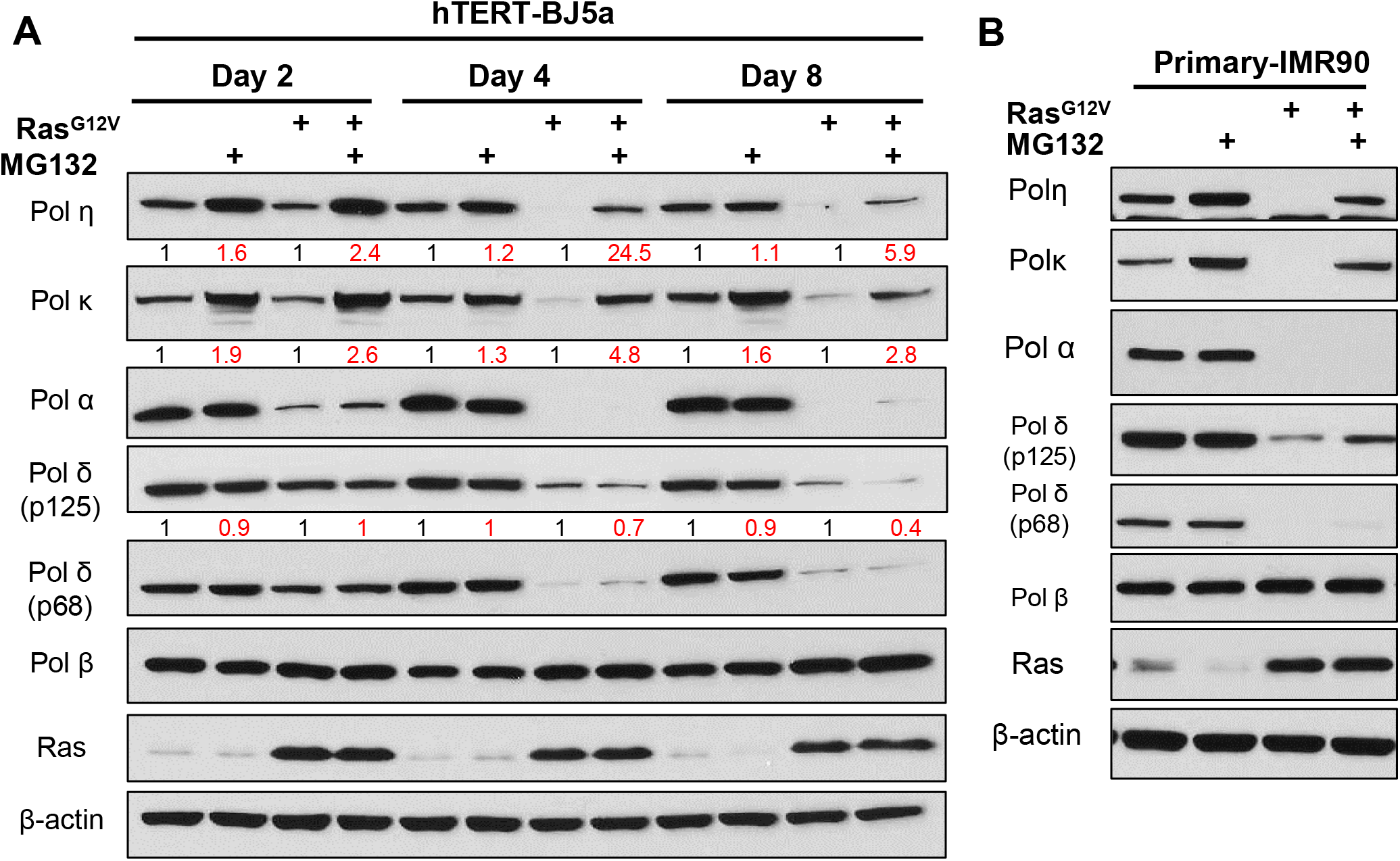
Ras^G12V^ overexpression-induced depletion of DNA polymerases is mediated by the proteasomal degradation pathway. (A) Immunoblot analyses of polymerase levels at indicated days after transduction with control or Ras^G12V^ vectors. BJ5a cells were either treated with DMSO or MG132 (10μM) for 4 hours prior to harvesting. Red values are quantification of polymerase levels, normalized to Control or Ras^G12V^ treated with DMSO. (B) Immunoblot analyses of primary fibroblast IMR90 cells infected with control or Ras^G12V^. On Day 8, cells were treated with DMSO or MG132 (10μM) for 4 hours prior to harvesting. One of two experiments is shown.

### DNA polymerase η depletion after Ras^G12V^ is partially mediated by MEK

We sought to determine if DNA polymerase downregulation is a direct result of the canonical Ras-MEK-ERK signaling pathway. Replicative DNA polymerase genes are regulated by E2F (reviewed in Barnes and Eckert 2017), and a tumor suppressor program known as senescence-associated protein degradation (SAPD) has been reported, in which certain proteins are selectively degraded in an ERK-dependent manner (Deschenes-Simard et al. 2013). Therefore, we hypothesized that inhibition of MEK-ERK signaling may restore both DNA polymerase gene expression and protein levels. To test this, we used the MEK specific inhibitor, Trametinib, to block ERK phosphorylation after Ras^G12V^ OE (Figure 3A-D). In control cells, we observed significant downregulation of POLA1, POLD1, POLD3, and POLE1 gene expression after 24 hours of Trametinib treatment (Figure 3B). This result is consistent with previous studies, and confirms that the MEK/ERK/E2F signaling axis is a positive regulator of replicative polymerase gene expression under normal cell culture conditions. We also measured a small but statistically significant increase in POLH gene expression after Trametinib treatment, suggesting that MEK signaling may negatively regulate POLH expression. Consistent with Figure 1, Ras^G12V^ OE significantly downregulated replicative polymerase gene expression. However, this repression was not alleviated upon MEK inhibition, and in several instances, Trametinib treatment of Ras^G12V^ OE cells further decreased POL gene expression (Figure 3A). Thus, Ras-induced downregulation of replicative polymerase gene expression is independent of the MEK-ERK signaling axis.

**Figure 3.**
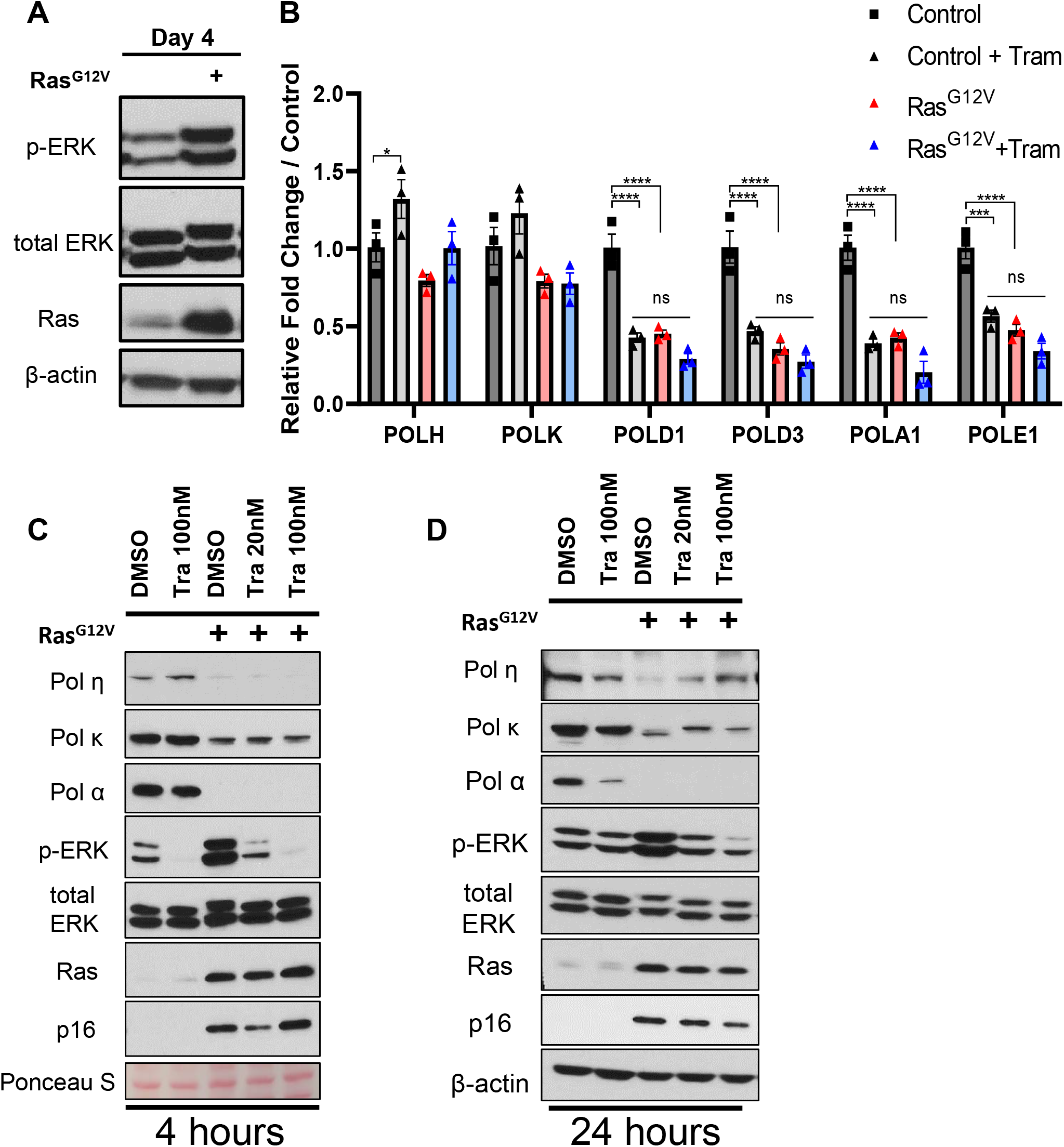
Ras^G12V^-induced depletion of DNA polymerase genes is MEK-independent, but Pol η depletion is partially dependent on ERK phosphorylation. (A) Immunoblot analysis of ERK phosphorylation in control and Ras^G12V^ transduced BJ5a cells at day 4. (B) Expression of Pol genes at Day 4 after infection of hTERT-BJ5a cells with control or Ras^G12V^ vectors. mRNA levels were determined using qRT-PCR after treatment of cells with Trametinib (100nM) for 24 hours prior to harvest. Data represent mean +/− SEM of three biological replicates. Statistical analyses were performed using Holm’s-Sidak multiple t-tests. *p<0.05, ***p<0.001, ****p<0.0001. n.s., not significant (C) Immunoblot analysis of control and Ras^G12V^ transduced BJ5a cells at Day 8. Cells were treated for 4 hours with DMSO or Trametinib (20nM, 100nM) prior to harvesting. Data are representative of two independent replicates. (D) Immunoblot analysis of control and Ras^G12V^ BJ5a cells at Day 8 and after 24 hours of treatment with DMSO or Trametinib (20nM, 100nM). Data are representative of three independent replicates.

Next, we treated Ras^G12V^ OE cells (8 days) with Trametinib for 4 or 24 hours to determine whether DNA polymerase protein depletion is mediated by MEK. We observed no rescue of Pols α and κ after Trametinib treatment and inhibition of ERK phosphorylation (Figure 3C-D). However, Pol η levels were partially restored after 24 hours of treatment, suggesting that depletion of DNA polymerase η is mediated, in part, by the MEK/ERK pathway. Overall, these data suggest that depletion of DNA polymerase gene and protein expression in response to Ras^G12V^ OE is primarily mediated by MEK-independent pathways.

### Senescence-associated p16^INK4A^ is not a major regulator of DNA polymerase depletion during Ras^G12V^ activation

We sought to ascertain whether the depletion of DNA polymerases is a general consequence of activating the cellular senescence program. We tested this using two approaches. First, we used the lentiviral transduction system to knockdown expression of RRM2 in BJ5a cells. Ras^G12V^ OE downregulates RRM2 expression, and loss of RRM2 is sufficient to induce senescence (Aird et al. 2013). As expected, RRM2 depletion induced markers of senescence, including increased p16, increased SA-β-galactosidase staining and decreased clonogenic survival (Figure 4—figure supplement 1A-B). We observed some decrease in DNA polymerase levels after 8 days of RRM2 knockdown (Figure 4—figure supplement 1B). However, the alterations were not nearly as dramatic as those measured in Ras^G12V^ OE cells, where depletion of Pols α, η, and κ was detected as early as day 2 (Figure 1B). For instance, although Pol α protein was modestly reduced 8 days after RRM2 knockdown, Pol α levels were at or below detection within 2 days after Ras^G12V^ OE (Figure 1A). Moreover, we did not observe a dramatic depletion of specialized Pols η and κ in the corresponding time frame for RRM2 knockdown cells (Figure 4—figure supplement 4B).

Second, we used an shRNA approach to knockdown CDKN2A (p16 ^INK4A^) expression after Ras^G12V^ (Figure 4—figure supplement 1C). The tumor suppressor gene CDKN2A (p16^INK4A^) is a major regulator of senescence (Serrano et al. 1997; Collado et al. 2005; Rayess et al. 2012), and we reasoned that depleting p16 levels might prevent polymerase downregulation. We observed a minor rescue of POLD1 gene expression but not any other polymerase genes after CDKN2A knockdown (Figure 4 A,B). At the protein level, we observed no statistically significant rescue of Pols α and κ at day 2 or day 8, when p16 ^INK4A^ levels were the lowest in Ras^G12V^ cells (Figure 4C,D). Unexpectedly, we discovered that Pol η protein expression is sensitive to the presence of p16 ^INK4A^. We observed a significant increase of Pol η levels at day 2, in both control and Ras^G12V^ cells with p16 knockdown. After 8 days of Ras^G12V^ OE, p16 levels are elevated even in the presence of lentiviral shCDKN2A; correspondingly, Pol η levels are depressed. Together, these results suggest that p16^INK4A^ negatively regulates Pol η; however, p16 is not the major regulator of DNA polymerase depletion in response to Ras^G12V^ signaling.

**Figure 4.**
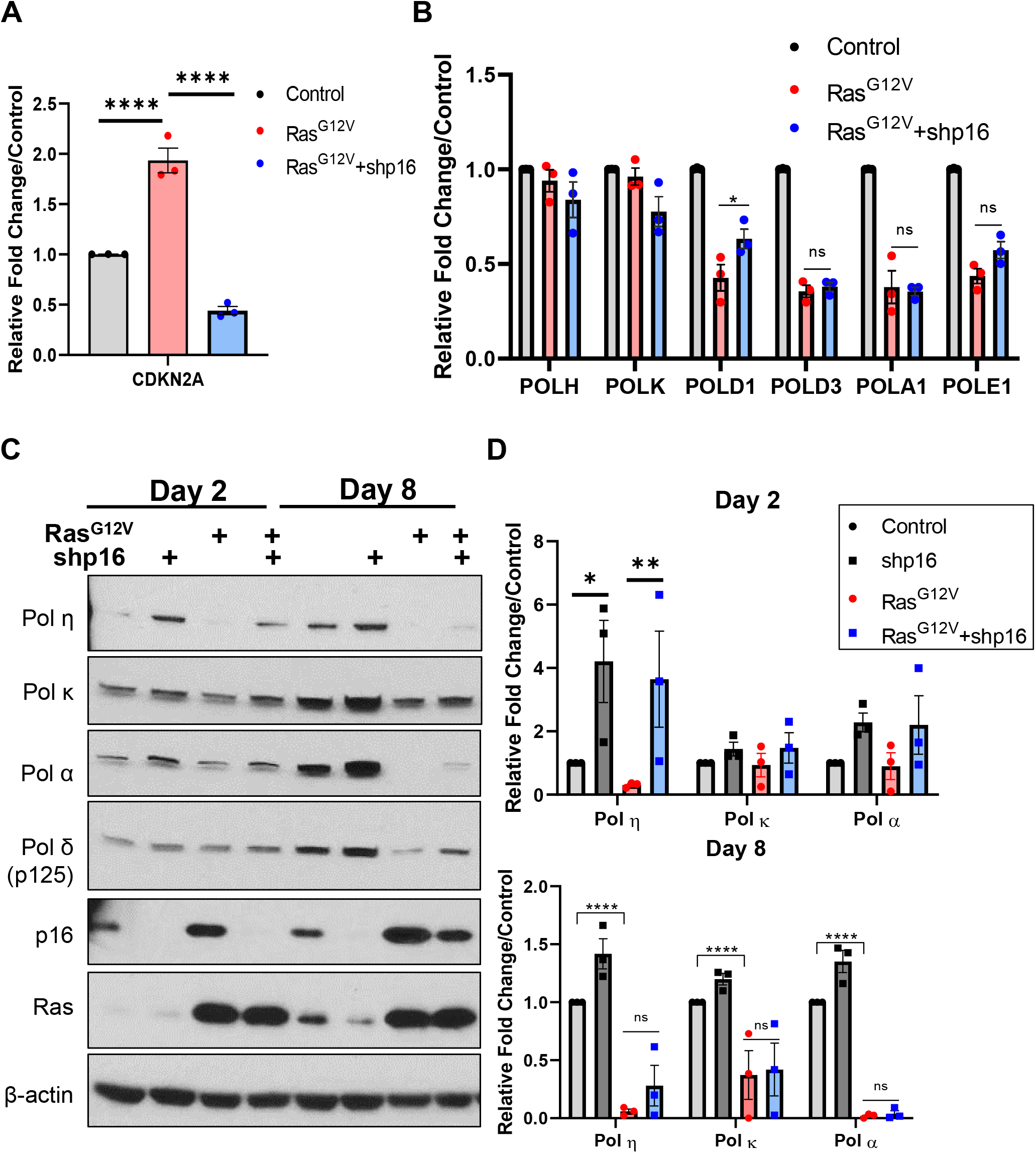
p16 is not a major regulator of DNA polymerase expression in response to oncogenic Ras^G12V^ signaling. (A) hTERT-BJ5a cells were infected with lentivirus expressing short hairpin RNAs (shRNAs) targeting CDKN2A. Scrambled shRNA was used as control. mRNA expression analysis of senescence markers on Day 2 as determined using qRT-PCR. Data represent mean +/− SEM from three biological replicates. ****p<0.0001. (B) mRNA analyses of DNA polymerase gene expression on Day 2 as determined using qRT-PCR. Data represent mean +/− SEM from three biological replicates. *p<0.05. (C) Immunoblot analyses of control or Ras^G12V^ BJ5a cells, with and without p16 knockdown. Data are representative of three independent replicates. (D) Quantification of control or Ras^G12V^ BJ5a cells immunoblot analyses, with and without p16 knockdown. Data represent mean +/− SD of three biological replicates. Statistical analyses were performed using Two-way ANOVA with Tukey’s post-hoc. *p<0.05, **p<0.01, ****p<0.0001. n.s., not significant.

### Pol η depletion induces a senescence-like growth arrest

The Ras-induced depletion of specialized Pols η and κ protein expression is intriguing, given that both polymerases play important roles in genome duplication, DNA damage, and the replication stress response (Sale et al. 2012; Bournique et al. 2018; Tonzi and Huang 2019). Specifically, we and others have shown that Pol η mRNA and protein levels are increased in tumor cells in response to exogenous sources of replication stress (Ceppi et al. 2009; Srivastava et al. 2015; Lerner et al. 2017; Barnes et al. 2018). To determine whether the presence of Pol η itself affects Ras^G12V^-mediated DNA polymerase depletion, we utilized hTERT-immortalized XPV cells without Pol η expression (hXPV), and isogenic POLH-complemented cells (hXPVη). After Ras^G12V^ OE, both hXPV and hXPVη cells displayed senescent phenotypes, including increased SA-β-galactosidase staining, decreased clonogenic survival, increased CDKN2A and decreased LAMINB1 mRNA expression, relative to controls (Figure 5—figure supplement 1A-C). Again, we observed robust depletion of DNA polymerases η, κ, and δ (p68 subunit) in both hXPV and hXPVη cells (Figure 5—figure supplement 1D). Moreover, the depletion of exogenous Pol η in hXPVη is further evidence that the regulation of Pol η is post-translational. These data show that Pol η expression is not necessary for Ras^G12V^-induced senescence and depletion of DNA polymerases in a non-transformed genetic background.

Previous reports have suggested that Pol η may play a role in regulating senescence. Pol η knockout mice display metabolic abnormalities and increased senescence-associated phenotypes specifically in adipocytes (Chen et al. 2015). Pol η also plays a role in suppressing senescence by participating in the alternative lengthening of telomeres (ALT) pathway (Garcia-Exposito et al. 2016). To test whether Pol η loss is sufficient to induce senescence, we transduced hTERT-BJ5a cells with lentiviral POLH shRNA. Strikingly, we found a significant reduction of Pol α protein expression and increased p16 expression (Figure 5A-C), concomitant with POLH knockdown. POLH depletion reduced the number of EdU positive cells on both days 2 and 8 (Figure 5D). POLH knockdown also increased SA-β-galactosidase staining and decreased clonogenic survival (Figure 5E-F). Taken together, these results suggest that depletion of Pol η decreases DNA polymerase α expression and is sufficient to induce a senescence-like growth arrest.

**Figure 5.**
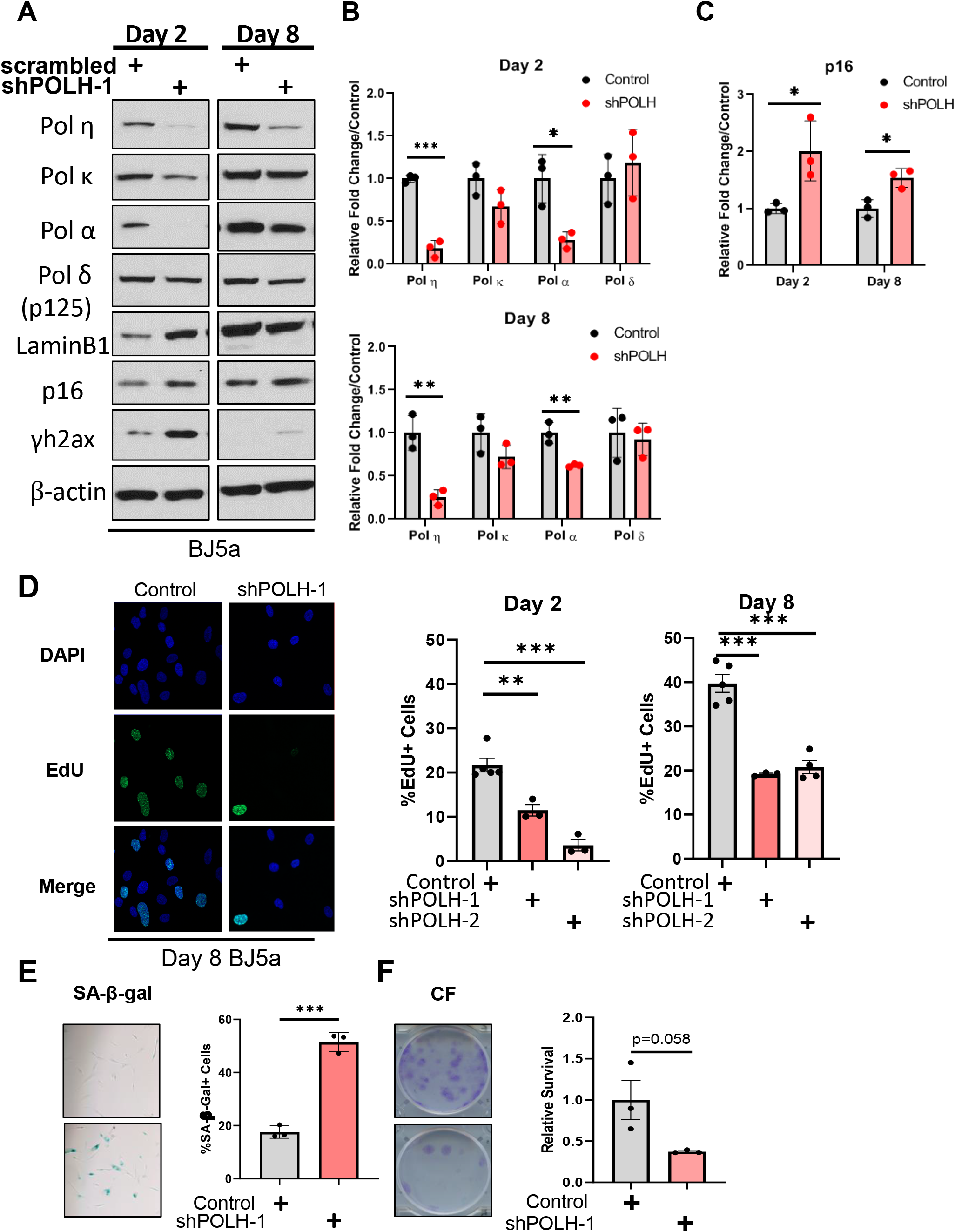
DNA polymerase η depletion is sufficient to induce a senescence-like growth arrest. (A) hTERT-BJ5a cells were infected with lentivirus expressing shRNA targeting POLH. Scrambled shRNA was used as control. Immunoblot analyses of indicated proteins at days 2 and 8 after lentivirus infection is representative of three biological replicates. (B) Quantification of immunoblot analyses of BJ5a cells infected POLH or scrambled shRNA lentivirus at days 2 and 8. Data represent mean +/− SD of three biological replicates. Statistical analyses were performed using Holm’s-Sidak method for multiple t-tests. *p<0.05, **p<0.01, ***p<0.001 (C) Quantification of p16 expression in BJ5a cells infected with POLH or scrambled shRNA lentivirus at days 2 and 8. Data represent mean +/− SD of three biological replicates. Statistical analyses were performed using Holm’s-Sidak method for multiple t-tests. *p<0.05. (D) Quantification of EdU positive cells (by immunofluorescence) of either two separate shRNA clones or scrambled shRNA at indicated days. Data represent mean +/− SEM for at least three biological replicates with two technical replicates. Statistical analyses were performed using Holm’s-Sidak method for multiple t-tests. **p<0.01, ***p<0.001 (E) Quantification of SA-β-gal activity at Day 8. Data represent mean +/− SEM. Statistical analysis was performed using unpaired Student’s t-test, two-tailed. ***p<0.001. (F) Quantification of clonogenic survival at Day 14. Data represents mean +/− SEM. Statistical analysis was performed using unpaired Student’s t-test, two-tailed.

### DNA polymerase degradation in response to Ras^G12V^ in SV40 transformed cells is associated with replication stress and improved survival

We and others have shown that the expression of DNA polymerases can be altered in human tumors (Lange et al. 2011); specifically, POLH is upregulated or amplified in melanoma, esophageal and ovarian cancer (Zhou et al. 2013; Tomicic et al. 2014; Srivastava et al. 2015; Tsao and Eckert 2018). To this end, we sought to determine whether the degradation of DNA polymerases after oncogene activation persists in transformed cells. To mimic a tumor-like model, we used immortalized, SV40 virus-transformed XPV30RO fibroblasts (SXPV) (Volpe and Cleaver 1995), in which the p53, Rb, and other cancer-associated signaling pathways are disrupted. To mimic the increased expression of Pol η observed in tumors, we utilized genetically related SV40 transformed, POLH-complemented XPV30RO cells (SXPVη), which overexpress Pol η (Kannouche et al. 2001). Similar to the hTERT and primary cell lines, Ras^G12V^ OE in SXPVη cells led to a rapid depletion of Pols α, η, κ and δ (p68 subunit) as early as day 2 and sustained through Day 8; however, depletion of the Pol δ catalytic p125 subunit was not observed (Figure 6A). In striking contrast, we observed no depletion of any DNA polymerase analyzed in SXPV (Pol η-deficient) cells after Ras^G12V^ OE. Importantly, our results show that DNA polymerase depletion in response to Ras^G12V^ occurs both in cells with a functional p53 (hTERT-BJ5a or hXPV, and IMR90 cells) and a non-functional p53 (SXPVη cells) pathway.

**Figure 6.**
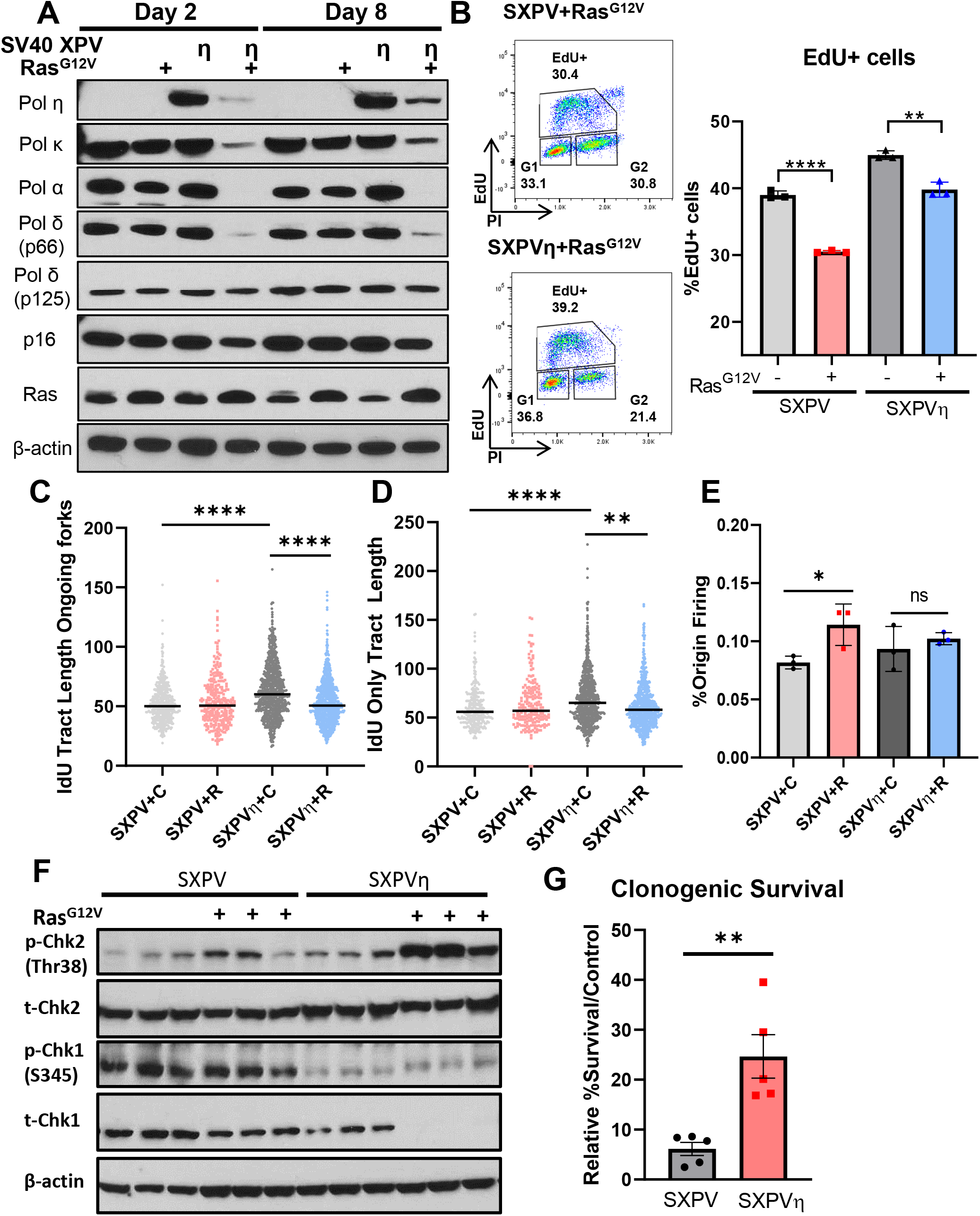
Oncogenic Ras^G12V^-induced DNA polymerase depletion is associated with fork slowing and altered checkpoint signaling. (A) Immunoblot analysis of SV40-transformed XPV cells (SXPV) and POLH-complemented SV40-transformed XPV cells (SXPVη), infected with control or Ras^G12V^ at Days 2 and 8. One of four biological replicates is shown. (B) *Left:* Representative cell cycle analyses of SXPV/SXPVη cells infected with Ras^G12V^ at Day 8. *Right:* Quantification of EdU positive cell population from three biological replicates. Statistical analysis was performed using unpaired Student’s t-test, two-tailed. **p<0.01, ****p<0.0001. (C) DNA fiber combing assay for ongoing (IdU-CldU) forks. Data represent IdU lengths of IdU-CldU tracts from two biological replicates. Statistical analysis was performed using Wilcoxon rank sum test with Holm’s post-hoc test. ***p<0.005, ****p<0.0001. (D) DNA fiber combing assay for terminated (IdU only) forks. Data represent IdU tract lengths from IdU only fiber of two biological replicates. Statistical analysis was performed using Wilcoxon rank sum test with Holm’s post-hoc test. **p<0.01, ****p<0.0001. (E) DNA fiber combing assay for origin firing. Percent origin firing was calculated using CldU only and CldU-IdU-CldU tracts / total tracts. Origin firing data is from three biological replicates. Statistical analysis was performed using unpaired Student’s t-test, two-tailed. *p<0.05. (F) Immunoblot analysis of control or Ras^G12V^-infected SXPV/SXPVη cells for indicated checkpoint proteins. All three biological replicates are shown. (G) Quantification of colony formation of Ras^G12V^-infected SXPV/SXPVη cells at Day 10, relative to respective controls. Data represent mean +/− SEM of five biological replicates. Statistical analysis was performed using unpaired Student’s t-test, two-tailed. **p<0.01.

Having this comparison of genetically related SV40 cell lines afforded us the opportunity to investigate the direct consequences of Ras-induced polymerase degradation. Replicative Pols α, δ, and ε are required for the bulk of on-going genome replication (Burgers and Kunkel 2017). Therefore, we asked what impact the severe depletion of the Pol α catalytic subunit has on DNA replication. Cell cycle analyses demonstrated that after 8 days of Ras^G12V^ OE, both SXPV and SXPVη populations contain ~30-40% of EdU+, S phase cells, a lower percentage than control infected cells (Figure 6B). Using DNA fiber analyses, we measured significantly longer replication fork tracts in control infected SXPVη compared to SXPV cells, suggesting that the presence of Pol η increases fork progression under unstressed conditions (Figure 6C). Typically, cells undergoing replication stress display slower fork progression, which can be compensated by increased origin firing (Kotsantis et al. 2018). Depletion of DNA polymerases in SXPVη after Ras^G12V^ OE led to a significant decrease in the length of ongoing and terminated forks, as expected for fork slowing (Figure 6C-D). However, we measured no increase in origin firing (Figure 6E), possibly due to low Pol α levels, a critical enzyme required for origin activation. In contrast, in Ras^G12V^-infected SXPV cells, where DNA polymerase depletion is absent, we observe no changes in fork progression. Instead, Ras^G12V^-infected SXPV cells displayed increased origin firing, compared to control infected cells (Figure 6D). These data demonstrate that the presence of Pol η in transformed cells alters replication dynamics and may be important for the induction of DNA polymerase depletion in response to Ras^G12V^ OE.

Oncogene-induced replication stress is well known to increase markers of DNA damage; therefore, we examined activation of the DNA damage response in both cell lines. In SXPVη cells with polymerase degradation, we observed a very robust increase in Thr68 phospho-Chk2 after 8 days of Ras^G12V^ OE, accompanied by a dramatic depletion of total Chk1 protein expression (Figure 6E). This observation is consistent with previous findings that cells suppress and deplete total Chk1 protein during genotoxic stress and nutrient deprivation via the senescence or the autophagy program (Zhang et al. 2005; Gabai et al. 2008; Kim et al. 2011; Park et al. 2015), perhaps as a mechanism of terminating the S phase checkpoint (Zhang et al. 2009). In contrast, after 8 days of Ras^G12V^ OE, SXPV cells without polymerase degradation showed no evidence of greatly increased Thr68 phospho-Chk2, while Chk1 was constitutively phosphorylated (Ser 345) (Figure 6E). Thus, polymerase depletion is correlated not only with altered fork progression, but also with differential checkpoint activation in response to oncogene overexpression. Finally, we examined the impact of replicative polymerase degradation on the ability of cells to survive prolonged oncogene overexpression. After 10 days of Ras^G12V^ OE, we measured a significantly greater clonogenic survival of SXPVη cells compared to SXPV cells (Figure 6F). Taken together, our results suggest that DNA polymerase depletion in response to Ras^G12V^ signaling is a protective mechanism, allowing cells to adapt to oncogene-induced replication stress by limiting replication fork progression and enforcing checkpoint activation, which ultimately improves cell survival.

## Discussion

DNA replication is a crucial phase of mitotic cell growth, as it represents the time during which the genome is most vulnerable. DNA polymerases are integral to all replication process, including ongoing fork elongation, fork restart, and fork repair. Despite their central role in replication, much remains unclear about the regulation of DNA polymerases in response to oncogene-induced stress and cellular senescence. In this study, we investigated the consequences of mutant H-Ras^G12V^ overexpression on DNA polymerase expression levels in human cells, focusing on a subset of polymerases known to be engaged at the replication fork. To our knowledge, our study is the first to report the robust cellular response to oncogenic Ras activation that invariably depletes multiple DNA polymerases. We discovered that distinct mechanisms regulate replicative *versus* specialized DNA polymerase expression: replicative polymerases are primarily transcriptionally repressed during Ras^G12V^ overexpression, while specialized polymerases are depleted via the proteasome degradation pathway. Inhibition of MEK signaling only partially rescued certain DNA polymerase levels, revealing that the mechanisms of DNA polymerase gene repression and protein degradation are both MEK-dependent and independent. Taken together, our study has uncovered a novel cellular response to oncogene-induced replication stress that is distinct from the response to exogenous replication stress, from the perspective of DNA polymerase regulation. This discovery has important implications for how human cells regulate DNA polymerases to limit DNA synthesis in response to endogenous replication stress, thereby limiting the detrimental consequences of oncogene expression on genome stability.

Oncogenic Ras^G12V^ overexpression in nontumorigenic cells induces an initial hyperproliferative response, followed by cellular senescence (Di Micco et al. 2006; Maya-Mendoza et al. 2015; Kotsantis et al. 2018). This response is accompanied by markers of replication stress, such as decreased fork elongation rate, and increased DNA damage, such as 53BP1 foci. Many mechanisms have been identified that contribute to oncogene-induced replication stress, including dysregulated replication initiation, altered nucleotide metabolism, transcriptional R-loops, and oxidative stress (Di Micco et al. 2006; Aguilera and Garcia-Muse 2012; Aird et al. 2013; Zeman and Cimprich 2014; Kotsantis et al. 2018). Here, we demonstrate a dramatic downregulation/degradation of DNA polymerases that are required for ongoing fork elongation, which we propose contributes directly to oncogene-induced replication stress. We discovered that the immortalized, SV-40 transformed XPV30RO cell line (Volpe and Cleaver 1995) fails to degrade DNA polymerases upon Ras^G12V^ OE; however, restoration of Pol η levels by POLH complementation (Kannouche et al. 2001) is sufficient to relieve the block on Ras^G12V^-induced DNA polymerase depletion (Figure 6). Using this experimental comparison, we demonstrate that cells undergoing oncogene-induced polymerase degradation display decreased replication fork progression, whereas fork lengths do not change in cells without DNA polymerase degradation (Figure 6). Surprisingly, even after 8 days of Ras^G12V^ OE, we observed that cell populations with reduced replicative polymerase levels retain EdU^+^, S-phase cells with slowed replication forks (Figure 6). This result is in concordance with a recent *in vitro* study of eukaryotic replisomes (Lewis et al. 2020). Using single molecule analyses and purified replication proteins, Lewis *et al.* show that the Pol α catalytic subunit was not required for continued elongation synthesis, suggesting that Pol α formation of initiator DNA from RNA primers is not an absolute requirement during lagging strand synthesis. In addition, a single Pol δ holoenzyme was retained in the replisome and could synthesize over 100 Okazaki fragments. We also measured robust Chk2 activation concomitant with Chk1 depletion in cells undergoing oncogene-induced polymerase degradation. Pol α and Pol κ both mediate ATR activation via the 9-1-1 complex (Yan and Michael 2009; Bétous et al. 2013), and we show that both polymerases are targets for Ras-induced depletion. These results suggest that the depletion of polymerases associated with the replication fork contributes to cell cycle checkpoint enforcement. Importantly, polymerase degradation likely is a protective response, as it is associated with improved cell survival in the face of oncogene-induced stress (Figure 6).

The coordinated degradation of replicative polymerases has been observed in fission yeast as a response to replication stress induced genetically in ΔSwi1 cells (ortholog of timeless) (Roseaulin et al. 2013). In that study, the forced accumulation of replication proteins was accompanied by excessive mitotic aberrations, suggesting that the degradation of replisome components plays a critical role in maintaining genome stability. We observed that Ras^G12V^ overexpression significantly down-regulated the Pol α catalytic subunit (POLA1 gene) and the Pol δ catalytic subunit (POLD1 gene). Extensive genetic studies using an *S. cerevisiae* model demonstrated that reduced expression of either Pol α or Pol δ induces substantially elevated rates of chromosomal loss and instability (Zheng and Petes 2018). The Pol δ accessory subunit, p68 (POLD3 gene) has a major effect on Pol δ’s PCNA binding affinity (Zhou et al. 2012) and mediates Pol δ’s retention in the replisome (Lewis et al. 2020). Our discovery that p68 (POLD3) also is a target for Ras-induced degradation is provocative, given that p68 is required for break-induced replication (Costantino et al. 2014; Dilley et al. 2016) and for mitosis-associated DNA synthesis (Minocherhomji et al. 2015).

The transcription factors E2F and Sp1 regulate polymerase gene expression under normal conditions (Barnes and Eckert 2017). Consistent with this, we show here that ERK positively regulates replicative polymerase genes under control conditions. We also show that basal POLH gene expression is distinct from replicative POL genes, as inhibition of MEK signaling increased rather than decreased POLH levels (Figure 3). Importantly, however, our study showed that the Ras^G12V^- mediated POL gene repression is MEK-independent (Figure 3). A Ras-induced senescence-associated protein degradation (SAPD) response has been reported, in which proteins are selectively degraded in an ERK and proteasome dependent process (Deschenes-Simard et al. 2013). Consistent with this report, we demonstrate that either MEK inhibition or proteasome inhibition partially rescued Pol η protein, suggesting that SAPD may be responsible for Ras^G12V^-induced Pol η depletion. Notably, however, these inhibitor treatments were not able to rescue replicative Pol α or δ proteins, supporting a predominantly transcriptional Ras^G12V^-induced repression program for replicative polymerases (Figures 2 and 3).

Pol η is heavily targeted for depletion and re-localization after DNA damage through several post-translational modifications, including phosphorylation, ubiquitination and SUMOylation (Bienko et al. 2010; Barkley et al. 2012; Despras et al. 2016; Bertoletti et al. 2017; Guerillon et al. 2020). We extend these previous findings by revealing a mechanism of Pol η depletion through activation of the MEK/ERK pathway of Ras signaling. Pol η also can be indirectly regulated via Rad18 phosphorylation through JNK signaling, another major pathway downstream of RAS (Barkley et al. 2012). The RAS family of proto-oncogenes (K-, H, N-Ras) mediate vital cellular processes such as growth, survival, metabolism, through several mitogenic pathways (Pylayeva-Gupta et al. 2011). Elucidating the exact mechanisms for transcriptional regulation and post-translational regulation of DNA polymerases downstream of the RAS signaling axes will significantly impact our understanding of the cellular replication stress response to oncogene activation.

Our discovery of Ras^G12V-^induced DNA polymerase depletion has significant implications for understanding the landscape of genome instability. Overexpression of oncogenes such as Ras^G12V^ and Cyclin E induce a unique landscape of common fragile site breakage (Miron et al. 2015). Fragile sites may contain difficult to replicate sequences (DiToRS), such as microsatellite sequences, non-B DNA structures, and R-loops, that are prone to double strand breaks (reviewed in Aguilera and Garcia-Muse 2012; Tsao and Eckert 2018). Specialized polymerases Pols η and κ are able to efficiently replicate DiToRS (Walsh et al. 2013; Barnes et al. 2017), and Pol η-deficient cells display elevated levels of replication stress and common fragile site breakage (Rey et al. 2009; Bergoglio et al. 2013; Barnes et al. 2018). Our findings here of Ras-induced Pol η and κ depletion raise the possibility that altered polymerase levels in cells undergoing oncogene-induced replication stress contribute to genome instability during neoplastic progression. One intriguing question is whether different oncogenes have differential impacts on polymerase expression levels, and whether this regulation is based on oncogene-induced replication stress intermediates, such as depleted nucleotides or unusual DNA secondary structures. For example, hydroxyurea induces replication stress by depleting nucleotide pools and requires Pol κ for replication stress tolerance (Tonzi et al. 2018). Recent studies have shown that Pol κ is preferentially utilized in Cyclin E/CDK2-mediated oncogenic stress (Yang et al. 2017), and the level of nuclear Pol κ is increased after treatment with BRAF, MEK, or ERK inhibitors (Temprine et al. 2020). In contrast, another replication stress inducer, aphidicolin, stalls DNA replication by inhibiting replicative polymerases and requires Pol η for tolerance (Rey et al. 2009; Barnes et al. 2018), and Pol η is utilized in Myc-mediated oncogenic stress (Kurashima et al. 2018). Together, current evidence supports the supposition that specific DNA polymerases are better suited for alleviating different causes of replication stress. Thus, elucidating the exact replication intermediates and signaling pathways that mediate DNA polymerase regulation, recruitment, and fork progression will be crucial to understanding the mechanisms underlying oncogene-induced replication stress and the roles of DNA polymerases in promoting tumorigenesis.

## Materials and Methods

### Cell Culture and Reagents

hTERT-immortalized BJ-5a human fibroblasts (CRL-4001™; ATCC) were cultured according to the ATCC guidelines in 4:1 mixture of Dulbecco’s medium and Medium 1999 supplemented with 10% Hyclone™ FBS (GE Healthcare) and 50μg/ml Gentimicin (Life Technologies). hTERT-immortalized XPV and POLH-complemented fibroblasts (gift from William Kauffman, University of North Carolina, Chapel Hill, USA) were cultured in Dulbeccos’ medium, 10% FBS, 50μg/ml Gentimicin, and 1% Minimal essential amino acids (Sigma). Experiments were performed on all cell lines between population doublings 10-35. SV40 transformed XPV and POLH complemented cell lines (XP30RO) were a gift from Jean-Sebastian Hoffman (Cancer Research Center. Toulouse, France) and were cultured in Dulbecco’s medium, 10% FBS, and 50μg/ml Gentimicin. Normal diploid IMR90 human fibroblasts (ATCC CCL-186) were cultured according to the ATCC in low oxygen (2% O_2_) in DMEM with 10% FBS supplemented with L-glutamine, non-essential amino acids, sodium pyruvate, and sodium bicarbonate. Experiments with IMR90 were performed between population doublings 25-35.

### Retro and lentiviral packaging and infection

Retrovirus production from pBabe vectors was performed using 293FT phoenix cells and human cell transduction was performed using the BBS/calcium chloride method (Aird et al. 2013). Lentivirus was packaged using the ViraPowerKit (Thermofisher Cat no. K497500) following the manufacturer’s instructions with modifications. Briefly, pLKO.1 Lentiviral constructs (Addgene) were transfected into 293FT cells using 35μg polyethylenimine per transfection (Alfa Aesar). After 48 hours of incubation, virus-containing media was collected. The experimental timeline of human cell infections and selection is outlined in Figure 1-figure supplement 1A. Cells were infected with viruses containing pBabe vector-only (control) or pBabe encoding HRas^G12V^, followed by a second round of infection 24 hours later. 24 hours after the second infection, the cells were replated, and selected with puromycin 6 hours after seeding (2μg/μl for BJ5a, 1μg/μl for hTERT and SV40 XPV cell lines). For dual Ras^G12V^ and shRNA lentiviral infections, BJ5a cells were first infected with viruses containing pBabe vector-only or HRas^G12V^ vectors. After 24 hours, a second round of infections was performed simultaneously with viruses containing both pBabe vectors and pLKO.1 vectors as described above. Cells were replated and selected with 4μg/ml puromycin for 2 days, followed by reseeding for subculture and assays. For infections with inhibitor treatments, infected cells were plated into 10cm^2^ plates the day before the indicated timepoints. After 24 hours, cells were treated with proteasome inhibitor MG132 (Sigma) or MEK inhibitor Trametinib (SelleckChem) for 4 hours or 24 hours, followed by RNA and protein isolation (described below).

### RT-qPCR

Experiments were performed according to MIQE guidelines with at least three technical replicates for all cell lines and three biological replicates for BJ5a cells. Total RNA was extracted using RNAeasy Kit (Qiagen), assessed for quality using the 2200 TapeStation (Agilent), and 600ng-1ug of samples with RIN>9 were converted to cDNA using qScript cDNA Synthesis Kit (Quanta Bioscences). qPCR was performed according to manufacturer guidelines with 20ng of cDNA, 1X Taqman target and control probes (Life Technologies), and PerfeCTa Fast Mix II, Low Rox (Quanta Biosciences). The reactions were analyzed using Agilent QuantStudio 7 Flex.

### Western blot and quantification

Whole cell extracts were collected by lysing cells with RIPA buffer (Santa Cruz Biotechnology) supplemented with Halt protease and phosphatase inhibitors (Life Technologies) and PMSF (Santa Cruz). Extracts concentrations were determined using DC™ Protein Assay (Bio-Rad). Sample preps were prepared using 4X LDS Sample buffer and 10X Reducing Agent (Life Technologies) and incubated at 70°C for 10min. Samples were loaded into pre-casted NuPage gels (Life Technologies). Gels were electrophoresed in 1X MOPS buffer, transferred onto 0.2um Amersham™ Hybond™ PVDF membranes (GE Healthcare). After transfer, efficiency was visualized by Ponceau S staining and blocked with 5% non-fat milk or BSA in TBS containing 0.1% Tween (TBST) before incubating with primary antibody overnight. Membranes were washed 5 times with TBST for at least 10min and were incubated with mouse or rabbit secondary antibodies at 1:20,000 dilution for 1 hour at room temperature. After washing with TBST for 5 times, blots were visualized with chemiluminescence reagents Amersham^TM^ ECL^TM^ Prime Western Blotting Detection Reagents (GE Healthcare) or Pierce™ ECL Western blotting substrate (Thermofisher). Bands were quantified using ImageJ (NIH) (Miller 2013) and normalized to β-actin.

### Beta-galactosidase staining

Senescence-associated (SA)-β-galactosidase staining was performed as previously described (Buj et al. 2019). Briefly, cells were fixed with 2% formaldehyde/0.2% glutaraldehyde in PBS for 10min. After washing with PBS, cells were stained at 37°C in a non-CO_2_ incubator with the staining solution (40mM Na_2_HPO_4_ pH 5.7, 150mM NaCl, 2mM MgCl_2_, 5mM K_3_Fe(CN)_6_, 5mM K_4_Fe(CN)_6_, 1mg/ml X-gal). After 16 hours, wells were washed with RO water and images were acquired using an inverted microscope (Nikon Eclipse Ti) with a 20X/0.45 objective and a camera (Nikon DS-Fi3).

### Immunoflourescence for EdU incorporation

Cells were plated on 22×22mm glass coverslips the day before indicated time point (Fisher Scientific). Cells were pulsed with 20μM EdU(5-ethynyl-2’deoxyuridine Lumiprobe) for 1 hour and fixed with 4% paraformaldehyde (Sigma) for 10min. Cells were washed with PBS and permeabilized with 0.3% Triton-X for 15min. Reaction mixes were freshly made with 20uM FAM-azide (Lumiprobe B5130), 4mM copper sulfate pentahydrate (Sigma), 20mg/ml ascorbic acid (Sigma) in PBS. Coverslips were labeled with 40μl reaction mix for 30min at room temperature. Coverslips were washed twice with PBS and mounted with antifade mounting medium VECTASHIELD with DAPI (Vectorlabs). Images were acquired at room temperature using Zeiss AXIO Microscope Imager.M2 and Apotome.2 apparatus with a 64X oil objective and the Zen Pro software. Representative pictures were obtained using Z-stacks and maximum intensity projections.

### Cell Cycle analysis

After infection, cells were fixed in 70% ethanol, washed, resuspended with 1% BSA in PBS, and placed in −20 °C for at least overnight until further processing. For standard cell cycle analysis, cells were resuspended with 0.1% Triton-X, 200μg/ml RNAase, and 40μg/ml propidium iodide. For EdU cell cycle analysis, cells were pulsed with 20μM EdU for 1 hour. Cells were harvested using trypsinization, fixed with 4% paraformaldehyde for 15min, and permeabilized with 0.1% saponin (47036-50G-F Sigma) and 1% BSA in PBS (Saponin-BSA buffer) for 15min. Cells were washed Saponin-BSA buffer and incubated with Click reaction buffer containing 20μM Sulfo-cyanin5 azide (Lumiprobe B3330), 4mM copper sulfate pentahydrate, and 20mg/ml ascorbic acide in PBS for 30min. Cells were washed with Saponin-BSA buffer twice and incubated with 200μg/ml RNAse and 40μg/ml propidium iodide at 37°C for 15min. Samples were analyzed on BD FACSCanto with FlowJo software. Cells with >4C DNA content were excluded from the analyses.

### Clonogenic Survival

Cells selected with puromycin for 2 days were seeded with triplicates in 6-well plates. Media was replaced every 2-3 days. After 10-14 days, cells were fixed with 100% methanol for 10min and stained with crystal violet solution for 15min. Wells were destained using 10% acetic acid and measured using 590nm absorbance with a spectrophotometer.

### Microfluidics-assisted replication track analysis (maRTA)

Fiber combing analyses were performed as previously described (Sidorova et al. 2009; Kehrli et al. 2016). Briefly, the day before the indicated timepoint, cells were plated in 60cm plates at 60% confluency overnight. Cells were labelled with 50μM IdU for 30min, washed 3 times with PBS, and labelled with 250μM CldU for 30min. Cells were trypsinized and washed with agarose insert buffer (10mM Tris7.5, 20mM NaCl, 50mM EDTA in water). After spin down, cells were resuspended in agarose insert buffer and mixed 1:1 with 2% low-melting agarose (Bio-Rad). Gel inserts were solidified at 4°C for overnight and were stored in agarose insert buffer until analysis. Microscopy of stretched DNAs was performed on the Zeiss Axiovert microscope with a 40x objective, and images were captured with the Zeiss AxioCam HRm camera. Fluorochromes were Alexa594 for CldU and Alexa488 for IdU. Lengths of tracks were measured manually in raw merged images using Zeiss AxioVision software, as well as automatically, using an open source software FiberQ (Ghesquiere et al. 2019) with concordant results. Percentages of ongoing (IdU-CldU) or terminated (IdU only) forks, and origin firing events were derived from FiberQ outputs. Statistical significance for track lengths was calculated using Kruskal-Wallis tests followed by pairwise Wilcoxon tests to derive p values adjusted for multiple comparisons. For origin firing percentages, statistical significance was determined in one-way ANOVA with post-hoc analysis with Tukey’s test to derive p values adjusted for multiple comparisons. Analyses were done in R Version 3.6.3.

### Statistics

Unless stated otherwise, graphical representation and statistical analyses were done using GraphPad Prism v8. Statistical analyses were performed as described in the figure legends.

## Acknowledgements

This study was supported by the National Institutes of Health (R01CA237153 to K.A.E., R00CA194309, R37CA240625 to K.M.A., R01GM115482 to J.M.S), by the Penn State Cancer Institute (Postdoctoral Fellowship to R.B.), and by the Jake Gittlen Memorial Cancer Research endowment. We thank Kelly Leon for expert technical assistance, and the Penn State Cancer Institute Flow Cytometry Core Facility. We are grateful to our colleagues for critical reading of our manuscript and useful suggestions for improvement.

## Author contributions

Conceptualization: WCT, KAE

Methodology: WCT, KMA, KAE, JMS, RB

Formal analysis: WCT, JMS, KAE

Investigation: WCT, JMS

Resources: KAE, KMA, JMS, RB

Writing—original draft: WCT

Writing—review & editing: WCT, RB, KMA, JMS, KAE

Visualization: WCT

Supervision: KMA, JMS, KAE

Project administration: KAE

Funding acquisition: KAE, JMS AND KMA

---

**Figure 1—figure supplement 1:**
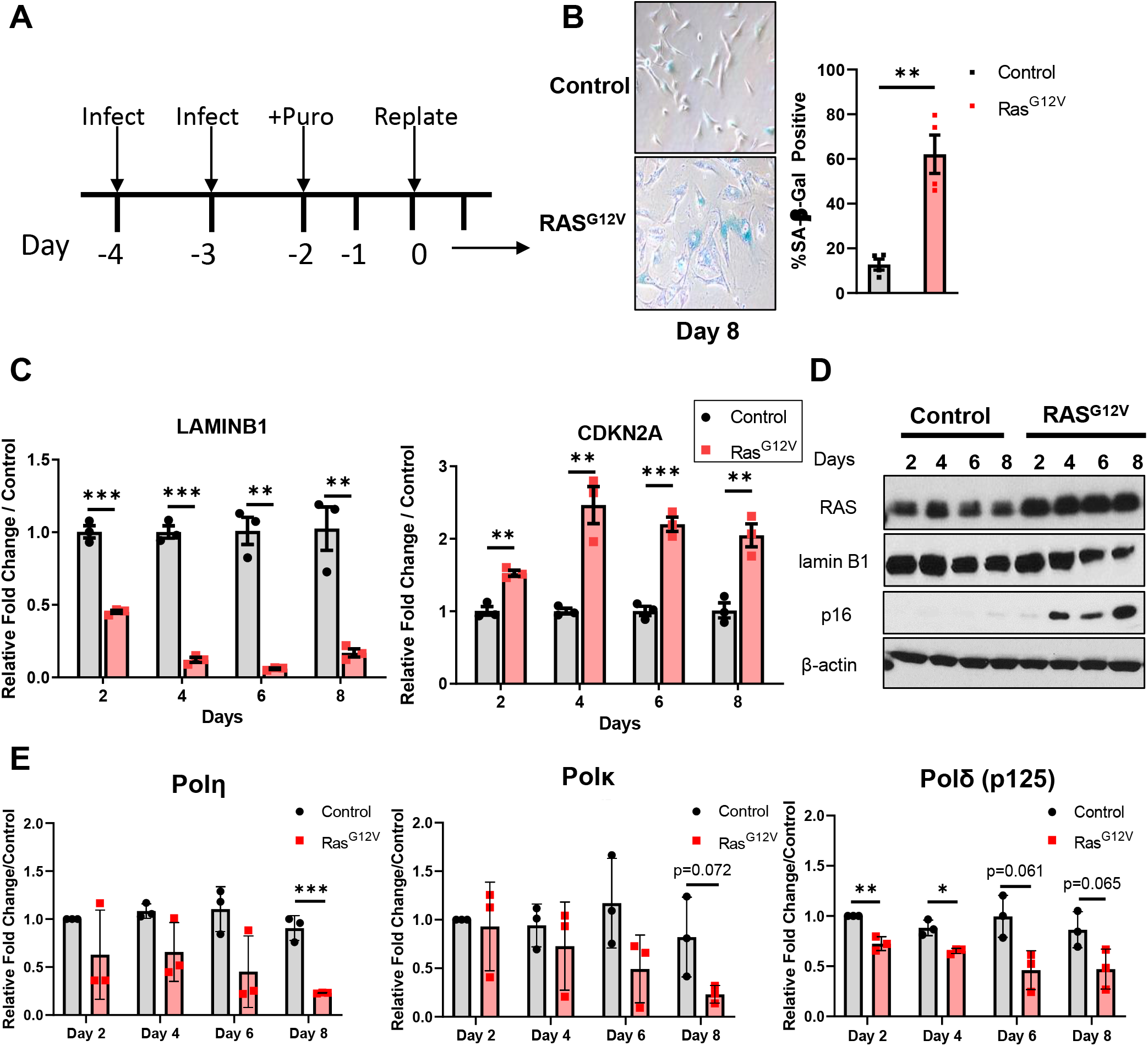
Ras^G12V^ overexpression in hTERT-BJ5a cells induces a senescent phenotype. (A) Schematic of the experimental infection and selection time course. (B) *Left panel:* SA-β-galactoside staining of BJ5a cells (Day 8) with and without Ras^G12V^ OE. *Right panel:* Quantification of β-galactosidase staining (N=3 technical replicates). Data represent mean +/− SEM. Statistical analysis was performed using Student’s unpaired t-test, two-tailed. **p<0.01. (C) mRNA expression of senescence markers Lamin B1 and CDKN2A. qRT-PCR was performed at the indicated timepoints. Data represent mean +/− SEM of three biological replicates. Statistical analysis was performed using Holm’s-Sidak multiple t-tests. **p<0.01, ***p<0.005. (D) Immunoblot analysis of senescence markers. One of three biological replicates is shown. (E) Time course of Pols η, κ, and δ protein depletion after Ras OE. Quantification of immunoblot analyses, at indicated times. Data represents mean +/− SEM three biological replicates. Statistical analyses were performed using Holm’s-Sidak multiple t-tests. *p<0.05, **p<0.01, ***p<0.005.

**Figure 1—figure supplement 2:**
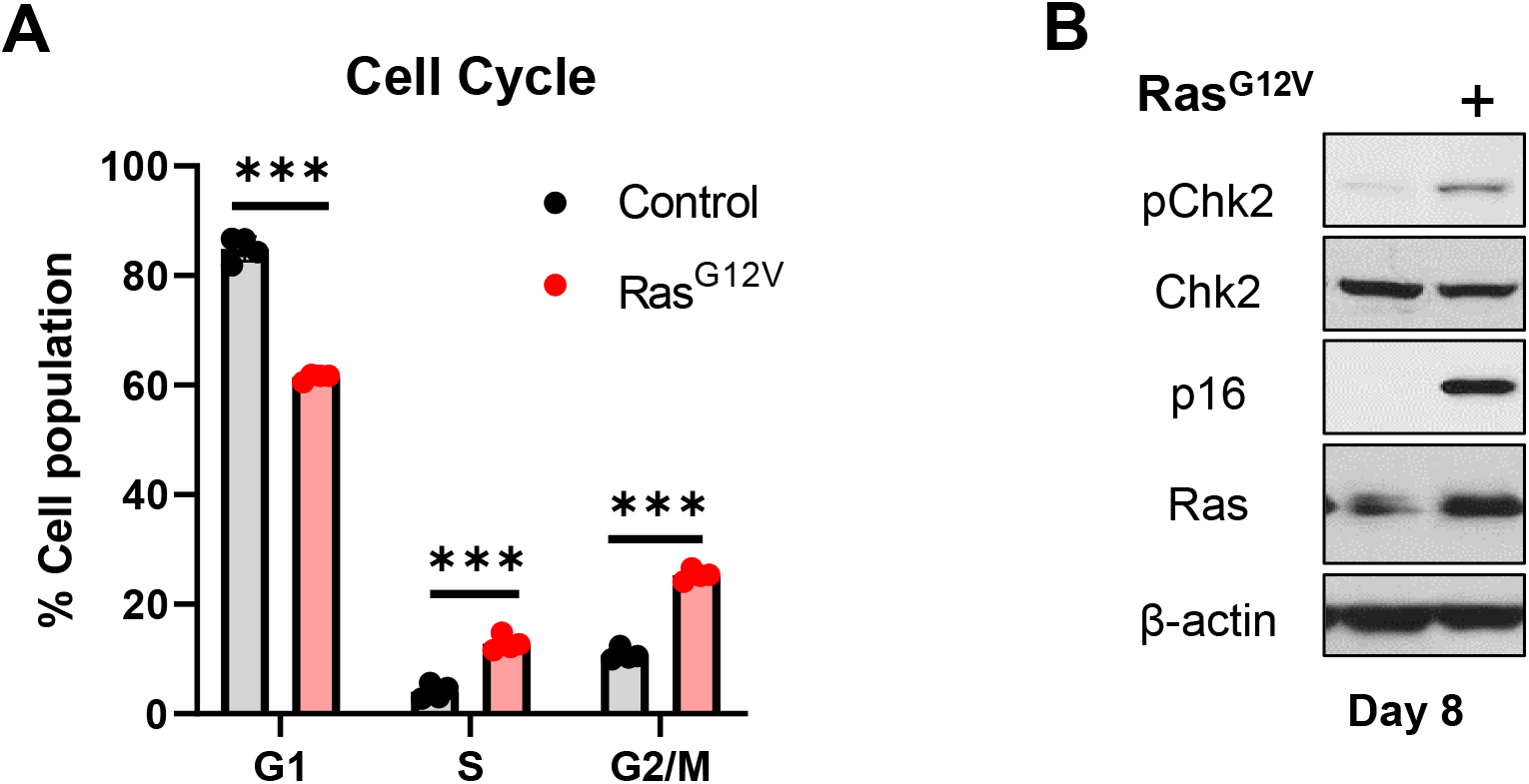
Cell cycle alteration and checkpoint response in hTERT-BJ5a cells transduced with Ras^G12V^. (A) Cell cycle (propidium iodide) analyses of control and Ras^G12V^ cells on Day 8. Data represent mean +/− SD of three biological replicates. Statistical analysis was performed using Holm’s-Sidak multiple t-tests. ***p<0.0001. (B) Checkpoint activation after Ras OE. Immunoblot analyses for Chk2 Thr38 phosphorylation on Day 8.

**Figure 2—figure supplement 1:**
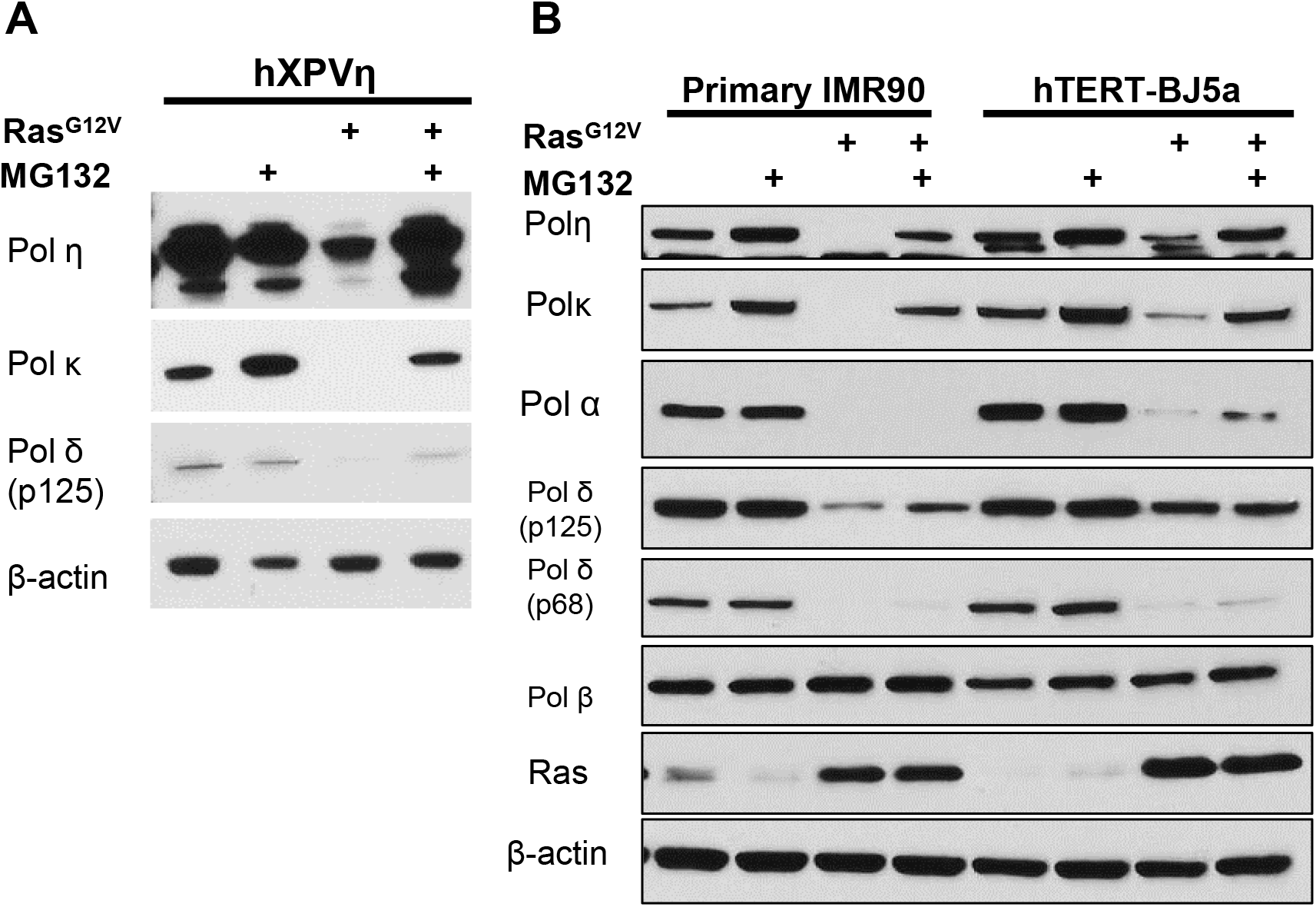
Ras^G12V^-induced depletion of DNA polymerases of DNA polymerases can be rescued in multiple fibroblast cell lines. (A) Immunoblot analysis of control or Ras^G12V^-infected hXPVη cells at day 8. Cells were treated with MG132 (10μM) for 4 hours prior to harvest. (B) Side-by-side comparison of IMR90 and BJ5a infected with control or Ras^G12V^. Cells were treated with DMSO or MG132 (10 μM) at day 8 for 4 hours prior to harvest. Note that the IMR90 results shown in this comparison are the same as those shown in Figure 2B, while the BJ5a results shown here are an independent biological replicate of Figure 2A.

**Figure 4—figure supplement 1:**
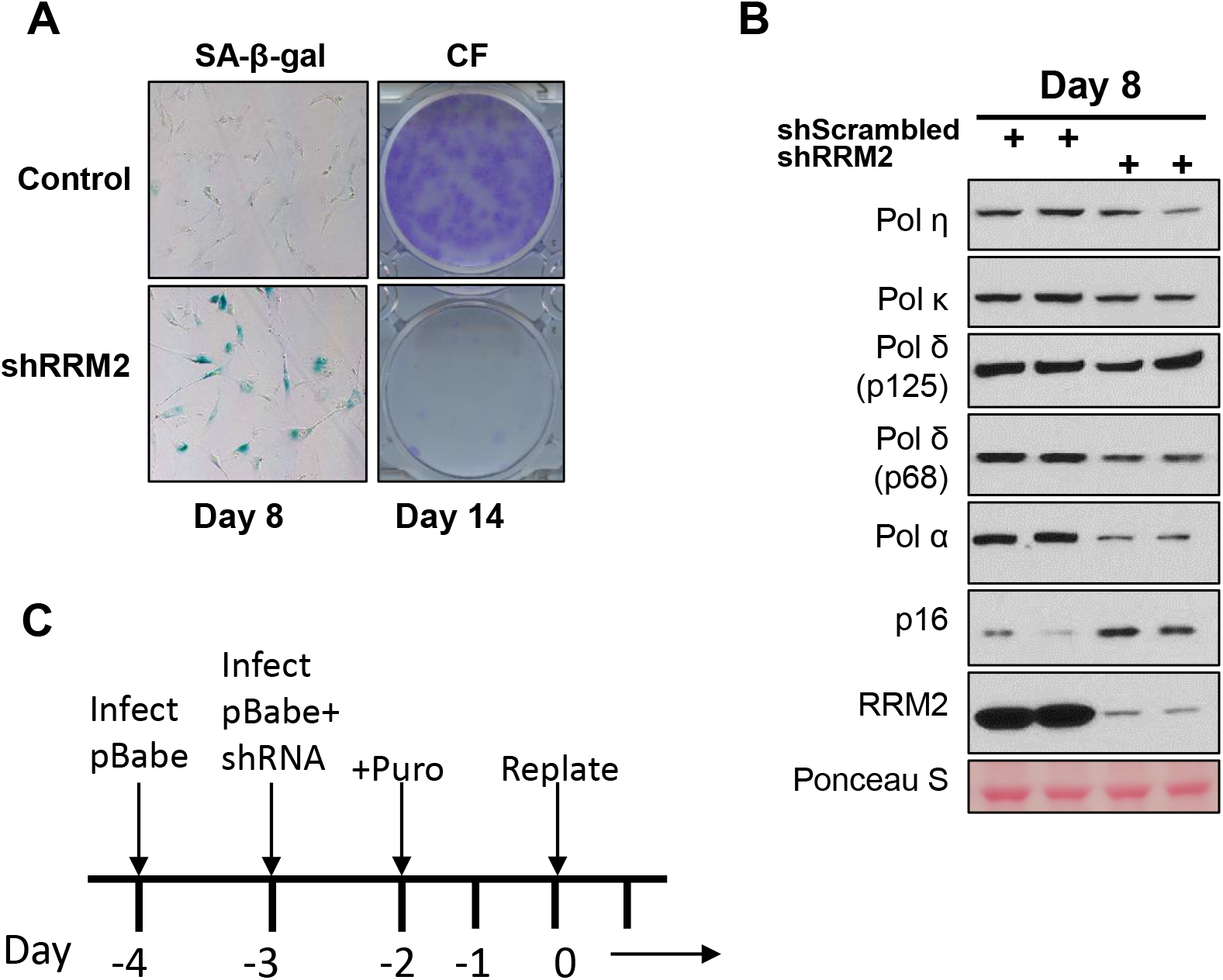
Rapid depletion of DNA polymerases is not apparent in shRRM2-mediated senescence. (A) hTERT-BJ5a cells were infected with lentivirus expressing short hairpin RNAs (shRNAs) targeting RRM2. Scrambled shRNA was used as control. SA-β-gal activity was assessed after Day 8 (Left). Colony formation of shRRM2 cells were assessed on Day 14 (Right). Results from one of two experiments is shown. (B) Immunoblot analyses of indicated proteins at Day 8 after lentivirus infection with control or shRRM2. (C) Schematic of Ras^G12V^ and shRNA knockdown approach.

**Figure 5—figure supplement 1:**
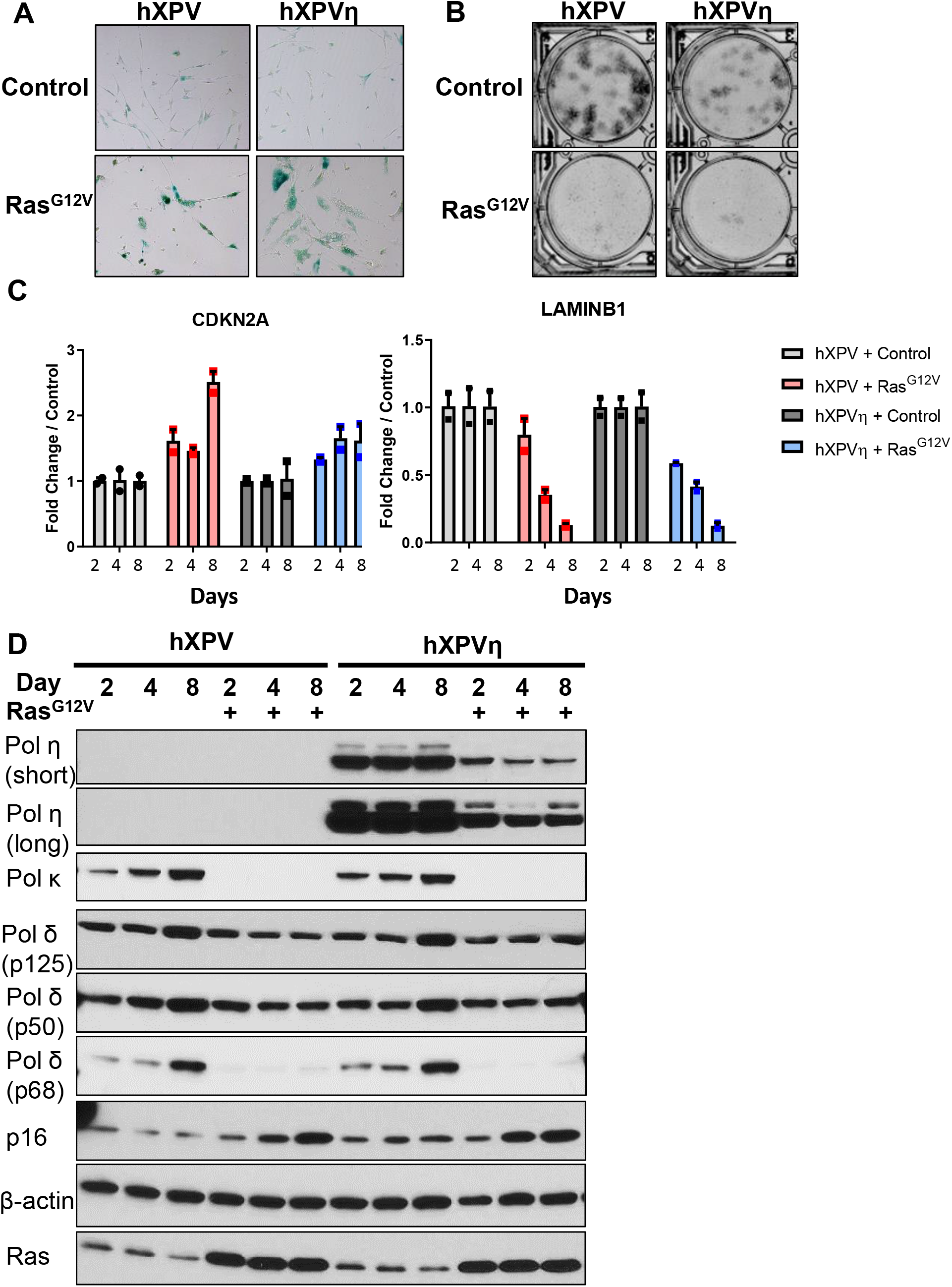
Ras^G12V^ induces oncogene-induced senescence response in human fibroblasts regardless of Pol η expression. (A) SA-β-galactosidase staining of hTERT-XPV (hXPV) and POLH complemented hTERT-XPV (hXPVη) cells with and without Ras^G12V^ OE (Day 8). One of two biological replicates is shown. (B) Colony formation assay of hXPV and hXPVη cells with and without Ras^G12V^ on Day 14. One of two biological replicates is shown. (C) CDKN2A and LAMINB1 mRNA expression in hXPV and hXPVη cells with and without Ras^G12V^ determined using qRT-PCR at the indicated days. Each gene was normalized to 18S internal control. Data represents mean +/− SD of two biological replicates. (D) Immunoblot analyses of hXPV and hXPVη fibroblasts with and without Ras^G12V^ at indicated days after selection. Data are representative of three biological replicates

